# Electrical stimulation of temporal and limbic circuitry produces distinct responses in human ventral temporal cortex

**DOI:** 10.1101/2022.07.06.498994

**Authors:** Harvey Huang, Nicholas M. Gregg, Gabriela Ojeda Valencia, Benjamin H. Brinkmann, Brian N. Lundstrom, Gregory A. Worrell, Kai J. Miller, Dora Hermes

## Abstract

The human ventral temporal cortex (VTC) is highly connected to integrate visual perceptual inputs with feedback from cognitive and emotional networks. In this study, we used electrical brain stimulation to understand how different inputs from multiple brain regions drive unique electrophysiological responses in the VTC.

We recorded intracranial EEG data in 6 patients implanted with intracranial electrodes for epilepsy surgery evaluation. Pairs of electrodes were stimulated with single pulse electrical stimulation, and corticocortical evoked potential (CCEP) responses were measured at electrodes in the collateral sulcus and lateral occipitotemporal sulcus of the VTC. Using a novel unsupervised machine learning method, we uncovered 2 to 4 distinct response shapes, termed basis profile curves (BPCs), at each recording electrode in the 11 to 500 ms post-stimulation interval.

CCEPs of unique shape and high amplitude were elicited following stimulation of several regions and classified into a set of four consensus BPCs across subjects. One of the consensus BPCs was primarily elicited by stimulation of the hippocampus; another by stimulation of the amygdala; a third by stimulation of lateral cortical sites, such as the middle temporal gyrus; and the final one by stimulation of multiple distributed sites. Stimulation also produced sustained high frequency power decreases and low frequency power increases that spanned multiple BPC categories.

Characterizing distinct shapes in stimulation responses provides a novel description of connectivity to the VTC and reveals significant differences in input from cortical and limbic structures.

**SIGNIFICANCE STATEMENT:** Disentangling the numerous input influences on highly connected areas in the brain is a critical step toward understanding how different brain networks work together to produce function. Single pulse electrical stimulation is an effective tool to accomplish this goal because the shapes and amplitudes of signals recorded from electrodes are informative of the synaptic physiology of the stimulation-driven inputs. We focused on targets in the ventral temporal cortex because it is an area strongly implicated in visual object perception. By using a data-driven clustering algorithm, we identified anatomical regions with distinct input connectivity profiles to the ventral temporal cortex. Examining high frequency power changes revealed possible modulation of excitability at the recording site induced by electrical stimulation of connected regions.

## INTRODUCTION

Distributed networks in the human brain, interacting with each other at multiple scales, have been described as the basis for our unique interactions with the sensory environment (Edelman and Mountcastle, 1978; Goldman-Rakic, 1988; Mesulam, 1990; Bressler, 1995; Sporns et al., 2004). Throughout the visual stream, feedforward processes from the stimulus-driven primary visual cortex converge with feedback influences from higher-level cognitive networks involved in attention, memory, and expectation (Moran and Desimone, 1985; Felleman and Van Essen, 1991; Motter, 1993; Schlack and Albright, 2007; McManus et al., 2011). The convergence of these processes in the ventral temporal cortex (VTC), for example, ultimately permits object recognition and the encoding of perceptual content, such as faces and scenery (Kanwisher et al., 1997; Epstein and Kanwisher, 1998). Functional connectivity between these networks has been explored primarily through correlated signals during behavioral tasks or a resting state, using a variety of electrophysiologic and imaging modalities (Friston, 1994; Kay and Yeatman, 2017). However, as multiple brain networks can be simultaneously active during a single task, conventional functional connectivity analysis limits the degree to which networks can be differentiated from one another.

An increasingly common method to characterize brain connectivity has been to measure evoked potentials in awake humans in response to single pulse electrical stimulation through implanted intracranial electrodes (Matsumoto et al., 2004; Lacruz et al., 2007; Kundu et al., 2020). Termed cortico-cortical evoked potentials (CCEPs), this technique enables the directional quantification of connectivity between any pair of implanted sites. In other words, it is a perturbational approach to determining effective connectivity. By measuring responses at one recording site of interest, we can calculate with high temporal resolution the influence of electrical stimulation inputs from stimulated sites sampled across different brain networks.

Earlier responses in CCEPs may represent signals propagated through direct pathways, while later responses may represent signals propagated through indirect or recurrent cortico-subcortico-cortical pathways (Matsumoto et al., 2004; Araki et al., 2015). Effectively, CCEPs can create a map of brain inputs to the recording site of interest.

Previous studies have quantified CCEPs through rigidly constrained metrics, such as amplitude or latency to negative or positive peaks within predetermined time intervals, e.g., the early “N1” peak (Matsumoto et al., 2004, 2007; Krieg, 2017; Kundu et al., 2020; Silverstein et al., 2020), but this approach often considers only the fastest, direct pathways and neglects the diversity of response shapes that are often present. A recent technique has captured this diversity by performing unsupervised clustering on CCEPs based on their variable temporal shapes (Miller et al., 2021). A unique canonical shape, termed basis profile curve (BPC), is calculated for CCEPs from each cluster of stimulation sites to the same recording site. Temporal deflections in intracranial EEG (iEEG) predominantly represent synchronous synaptic activity at the recording site (Mitzdorf and Singer, 1978). Since the recording site is constant for all CCEPs, different temporal motifs across BPCs may indicate different cortical layers in the same ensemble of neurons targeted by CCEP inputs, or they may indicate inputs to different ensembles of neurons altogether in the neighborhood of the recording electrode. Uncovering these input differences has significant functional and anatomical implications. For instance, the laminar patterns of neuronal inputs, as assessed by injected tracers, have formed the basis for determining feedforward and feedback connectivity within the visual system (Felleman and Van Essen, 1991; Markov et al., 2014). We use the BPC method here as a means of uncovering the signatures of such afferent inputs in awake human participants.

As the human VTC simultaneously receives inputs from many networks to accomplish its goal of visual perception, it serves as an optimal candidate for disambiguating network inputs by single pulse electrical stimulation. We have recorded CCEPs at electrodes in the VTC to parse input connectivity from independently stimulated sites across multiple brain networks. By applying the BPC method, we separate those inputs into distinct physiologically relevant categories informed by temporal shape.

## METHODS

### Subjects

IEEG voltage data were measured in 6 human subjects (4 female, see Table 1) who had been implanted with stereo electroencephalography (sEEG) electrodes for epilepsy monitoring and had electrodes placed in the VTC. Recorded data were filtered between 0.01 Hz and 878 Hz and then digitized at 2048 Hz on a Natus Quantum amplifier.

**Table 1.** Subject demographics. The table lists age, sex, implanted hemisphere, number of stimulation sites, and electrode IDs for each subject. The electrodes were enumerated from 1 starting in the collateral sulcus in each subject.

| Subject ID | Age | Sex | Hemisphere implanted | Collateral sulcus electrode ID | Lateral occipito-temporal sulcus electrode ID | Number of stimulation sites |
| --- | --- | --- | --- | --- | --- | --- |
| 1 | 13 | F | L | 1, 2, 3, 4 |  | 41 |
| 2 | 30 | F | R | 1 |  | 52 |
| 3 | 29 | F | L | 1 |  | 31 |
| 4 | 20 | F | L | 1, 2, 3, 4, 5 |  | 45 |
| 5 | 46 | M | R | 1, 2, 3 | 4, 5 | 42 |
| 6 | 18 | M | R |  | 1, 2 | 36 |

### Electrode localization

Subject preoperative T1 MRIs were first transformed into AC-PC space through affine transformations and trilinear voxel interpolation (Huang et al., 2021). sEEG electrodes were then localized from the postoperative CT scans and co-registered to the T1 MRIs (Hermes et al., 2010), so that electrode positions were in AC-PC space.

Subject T1 MRIs were segmented using the autosegmentation algorithm in Freesurfer 7 (Dale et al., 1999), which also produced a 3D mesh rendering of the pial surface. Gyral and sulcal labels were generated for each subject’s pial surface, during autosegmentation, by aligning surface topology to the Destrieux cortical atlas (Destrieux et al., 2010). Electrodes were matched to cortical or subcortical labels corresponding to the most frequent voxel label within a 3 mm radius, and these labels were visually reviewed for correct assignment. Electrodes were visualized in AC-PC space on individual subject T1 MRI slices or on subject pial and inflated pial surfaces.

For visualization purposes, electrode positions were also transformed to the standard Montreal Neurological Institute (MNI) 152 space using non-linear segmentation-based normalization of the T1 scan in SPM12 (Penny et al., 2011) (https://www.fil.ion.ucl.ac.uk/spm/), so that recording and stimulation sites across subjects can be visualized on the same MNI 152 pial surface rendering and MNI 152 T1 MRI slices. Positions in the right hemisphere were reflected across the mid-sagittal (yz) plane when rendered to the MNI 152 pial surface, so that all positions could be visualized on a single (left) hemispheric rendering.

### Electrode of interest selection

After preprocessing, we studied inputs to electrodes in the collateral sulcus (Destrieux label *S_oc-temp_med_and_Lingual*) and the lateral occipito-temporal sulcus (Destrieux label *S_oc-temp_lat*). These regions were implanted in multiple subjects and allowed us to test the hypothesis that different inputs cluster systematically across subjects.

### Experimental Design and Preprocessing

Electrode pairs were stimulated up to 36 times each (average 10) with a single biphasic pulse of 200 microseconds pulse width and either 4 mA or 6 mA amplitude every 3-7 seconds, using a Nicolet Cortical Stimulator. Electrodes and individual trials were visually inspected, and those that showed artifacts or epileptiform activity were excluded from analysis. Stimulation sites with fewer than 8 remaining stimulation trials were also excluded from analysis. Table 1 shows the final number of stimulation sites analyzed in each subject. Note that this number includes stimulation sites that were excluded in the analysis of individual recording electrodes, when the stimulation pair contained the recording electrode. Data were structured to conform to the iEEG-BIDS format (Holdgraf et al., 2019).

All preprocessing was performed using custom Matlab code (see Code Availability, below). Data in all subjects were re-referenced on a trial-by-trial basis to a modified common average that avoids introducing large stimulation effects or evoked potentials in other electrodes. Specifically, the reference was the mean of all channels that were not stimulated, had below 95 percentile variance between 500-1000 ms post-stimulation, and whose variance between 15-100 ms post-stimulation was less than 1.5-fold that between 500-1000 ms post-stimulation. Line noise at 60 Hz and the first two harmonics were then removed using the spectral interpolation method implemented in (Mewett et al., 2001). Finally, the median baseline voltage value from 500-50 ms pre-stimulation was subtracted from each individual trial and channel.

### Subject-level BPCs

For each recording site in each subject, BPCs were calculated from the single-pulse CCEP data based on the method described in Miller et al. (2021). Each BPC is a single unit-normalized curve that characterizes the temporal shape common to CCEPs from a set of stimulation sites to the recording site. It is also possible for some stimulation sites (usually those with weak CCEP responses) to be not assigned to any BPC. We applied the algorithm with a few minor modifications, as follows (Figure 1). First, the BPC method takes as input a matrix of stimulation trials recorded at one electrode, in this case in the collateral sulcus or lateral occipito-temporal sulcus (Figure 1A). In our implementation, each trial was weighted by an exponential decay function with a time constant of 100 ms, in order to bias the BPC algorithm in favor of earlier responses (Figure 1B). Second, BPCs were calculated on the 11-500 ms interval after electrical stimulation. The 11 ms start time was determined to be the earliest time point without the influence of stimulus artifacts. Pairwise scalar projections were calculated between all CCEP trials and t-statistics were calculated from the projection values, grouped by stimulation site pairs, to generate a significance matrix, Ξ. This significance matrix quantified the temporal similarity between inputs from pairs of stimulation sites (e.g., CCEPs from stimulation sites 1 and 2 in Figure 1A are highly similar to each other and thus had a high t-statistic in Ξ). Then, non-negative matrix factorization (NNMF) was performed on Ξ to yield components W and H, such that Ξ ∼ WH. NNMF is necessary to ensure that temporal dynamics from each cluster do not contribute both positively and negatively to CCEPs recorded at the same electrode (i.e., laminar anatomy is not invertible). In our implementation, the NNMF convergence variable, ζ, was defined as the *sum* of the upper-half off-diagonal elements in HH^T^, and the convergence threshold for ζ was set at 1. This resulted in a low-rank decomposition of the CCEP signals. A winner-takes-all decision then assigned each stimulation site either to one or no BPC group, based on its coefficient in H (no BPC group is assigned if the minimum threshold is not met for that stimulation site, justification in Miller et al., 2021). BPC shapes were then determined as the first principal component of CCEPs from each NNMF group of stimulation sites, and they were visualized after multiplication by the reciprocal of the exponential weighting function (Figure 1C). For example, stimulation sites 1 and 2 cluster into the third BPC (green) shown in Figure 1C. If a BPC is assigned, a numerical signal-to-noise ratio (SNR), quantifying the fit of the BPC relative to residual noise, is also calculated for each trial at that stimulation site:

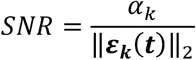

where the k^th^ CCEP trial ***Vk*(*t*)** = *αk* ***B*(*t*) + *εk*(*t*)**, and ***B*(*t*)** is the BPC assigned for that trial. The mean SNR across trials is used as the average BPC fit for each stimulation site.

**Figure 1.**
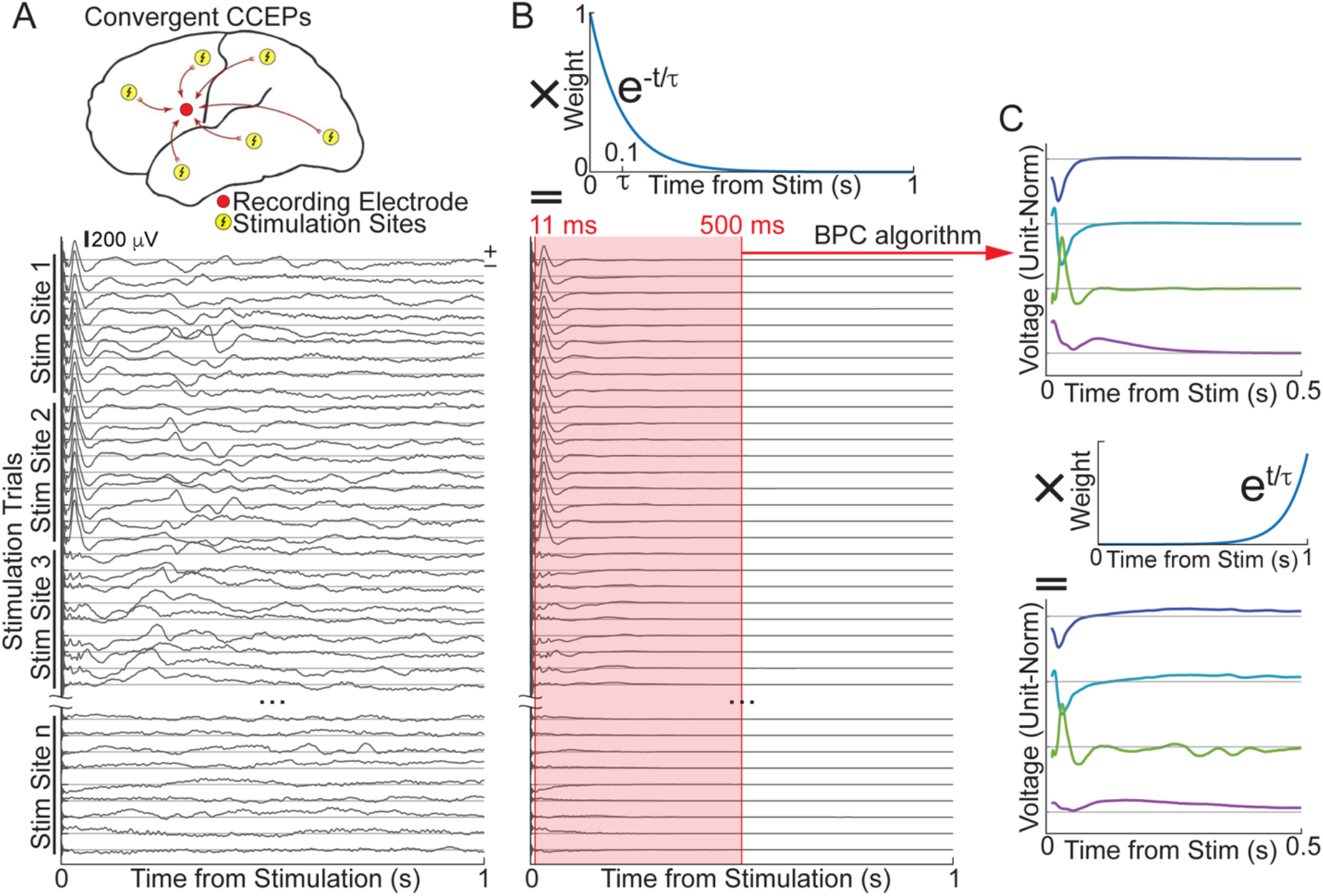
Modified procedure for identifying basis profile curves (BPCs) from CCEP data. **A**, Voltage data (positive is up) were recorded in a convergent paradigm from a fixed electrode while multiple electrode pairs were stimulated (top) to yield input CCEP data (bottom). Multiple stimulation trials (average = 10) were performed at each stimulation site. Top illustration was adapted from Miller et al., 2021. **B**, Each CCEP trial was multiplied by an exponential decay weighting function with time constant, τ, equal to 100 ms. All CCEPs from 11 to 500 ms post-stimulation were used to calculate BPCs. **C**, BPCs were multiplied by the reciprocal of the exponential weighting function in B for visualization.

### Group level consensus BPCs

We calculated a single set of “consensus” BPCs across all collateral sulcus electrodes in subjects 1 through 5, as follows. All BPCs across electrodes and subjects were concatenated to form a matrix, *A*, where columns correspond to BPCs and rows correspond to time points. Singular value decomposition was applied to *A*, to yield orthogonal eigenvectors of *AA*^*T*^ in descending order of variance explained. This equated to principal component analysis without centering on features (time points). The first 3 eigenvectors of *AA*^*T*^ formed a low-dimensional basis on which subject BPCs were clustered using K-means clustering. K=4 clusters were chosen based on visual inspection of the subject BPCs projected to the first 3 eigenvectors. The consensus BPCs were obtained by projecting the centroid of each cluster back into signal space and labeled arbitrarily in increasing order by coefficient along the first principal component. Each subject BPC was assigned to a consensus BPC based on its K-means cluster. This ultimately yielded consensus BPC labels for all stimulation sites that possessed subject-level BPC labels, across all input electrodes and subjects.

For each consensus BPC, we tallied the number of stimulation sites that were located in each of six large cortical or subcortical regions. Stimulation sites whose SNR were less than 1 were excluded from this analysis. First, each stimulation site was matched to an anatomical label in a similar way as for electrodes, with the position of each stimulation site being defined as the midpoint position between the pair of stimulated electrodes. Next, gyral and sulcal labels (based on the Destrieux atlas) were binned together into six regions as follows. The ventral temporal region included the fusiform gyrus (*G_oc-temp_lat-fusifor*), lingual gyrus *(G_oc-temp_med-Lingual*), parahippocampal gyrus *(G_oc-temp_med-parahip*), lateral occipito-temporal sulcus (*S_oc-temp_lat*), collateral sulcus (*S_oc-temp_med_and_Lingual*), anterior collateral sulcus (*S_collat_transv_ant*), and posterior collateral sulcus (*S_collat_transv_post*). The lateral temporal region included the inferior temporal gyrus (*G_temporal_inf*), middle temporal gyrus (*G_temporal_middle*), superior temporal gyrus (*G_temp_sup-Lateral, G_temp_sup-Plan_polar, G_temp_sup-Plan_tempo*) temporal pole (*Pole_temporal*), Heschl’s gyrus (*G_temp_sup-G_T_transv*), inferior temporal sulcus (*S_temporal_inf*), superior temporal sulcus (*S_temporal_sup*), and transverse temporal sulcus (*S_temporal_transverse*). The insula included the long insular gyrus and central sulcus of the insula (*G_Ins_Ig_and_S_cent_ins*); short insular gyri (*G_insular_short*); and circular sulcus of the insula *(S_circular_insula_ant, S_circular_insula_inf, S_circular_insula_sup*). The hippocampus and amygdala were each their own region, and the “other” category included all other anatomical labels. Independence between region and consensus BPC across all stimulation sites (with robust SNR ≥ 1) was determined by using the Chi-Square Test of Independence.

Consensus BPC analysis was not performed on CCEP inputs to the 4 lateral occipito-temporal sulcus electrodes in subjects 5 and 6, due to the small number of recording electrodes and subjects (see Table 1).

### Spectral and broadband analysis

Induced wavelet spectrograms were calculated for each CCEP trial using the MATLAB cwt package on the preprocessed time series data. Power was calculated for each time-frequency bin as the square of the amplitude of the wavelet transform. Power was then normalized to baseline separately for each frequency by dividing by the mean power between 500-50 ms before stimulation. An average spectrogram was determined for each stimulation site by taking the geometric mean across trials. A single heatmap for each consensus BPC was then obtained by collecting the average log spectrograms from all stimulation sites assigned to that consensus BPC and calculating the one-sample t-statistic at each frequency-time bin against the null hypothesis of 0 (no log-fold change over baseline). As in the anatomical analysis, stimulation sites whose SNR were less than 1 were excluded from this analysis. A t-statistic heatmap was also calculated from all stimulation sites that were algorithmically excluded from BPC assignment, as a negative control.

A time-varying broadband estimate was also calculated for each CCEP trial, as follows. For each recording site in each subject, the preprocessed signals were band-pass filtered by forward-reverse filtering using a 3rd order Butterworth filter between 70 and 170 Hz. Subsequently, the broadband estimate was calculated as the log squared absolute value (log power) of the Hilbert Transform of each band-pass filtered signal. Broadband estimates were averaged across trials for each stimulation site, and then averaged again across all stimulation sites assigned to a given consensus BPC to yield a single time-varying broadband estimate for each consensus BPC. An average broadband estimate was also calculated from all stimulation sites that were algorithmically excluded from BPC assignment, as a negative control.

### Statistical analysis

Time-frequency bins significantly different from rest in the consensus BPC t-statistic heatmaps and samples significantly different from rest in the consensus BPC broadband estimates, by one-sample t-tests, were controlled for multiple comparisons by using a 5% false discovery rate under dependency, as implemented by (Benjamini and Yekutieli, 2001). This method was chosen as nearby time-frequency bins often exhibit positive or negative regression dependency with each other.

### Data and Code Accessibility

Data will be made available in BIDS on OpenNeuro and code will be made available on GitHub.

## RESULTS

To characterize distinct types of input connectivity to the ventral temporal cortex, we identified canonical waveforms using the BPC method on CCEP data recorded from electrodes in the collateral sulcus and lateral occipito-temporal sulcus. We calculated a single set of “consensus BPCs” across subjects and identified the anatomical distribution of stimulation sites that were assigned to each BPC. We also investigated the induced spectral power in each consensus BPC to understand the relationship between local neuronal activity and waveform shape.

### BPCs from collateral sulcus input CCEPs in each subject

Collateral sulcus inputs were recorded from five subjects (1-5 electrodes each) while stimulating between 31 and 52 electrode pairs (Table 1). Application of the BPC algorithm uncovered sets of 2 - 4 subject-level BPCs at each measurement electrode, representing distinct canonical CCEP shapes between 11 to 500 ms post-stimulation.

BPCs from electrode 1 in subject 1 are presented in Figure 2. Here, we observed three distinct shapes (Figure 2B). The first BPC was characterized by a prominent negative deflection at 27 ms, followed by a wide positive peak centered at 255 ms; the second BPC was characterized by an initial negative deflection at 31 ms, followed by two positive peaks centered at 81 ms and 327 ms; the third BPC was characterized by a first positive peak at 27 ms and gradual tapering to baseline over the entire duration.

**Figure 2.**
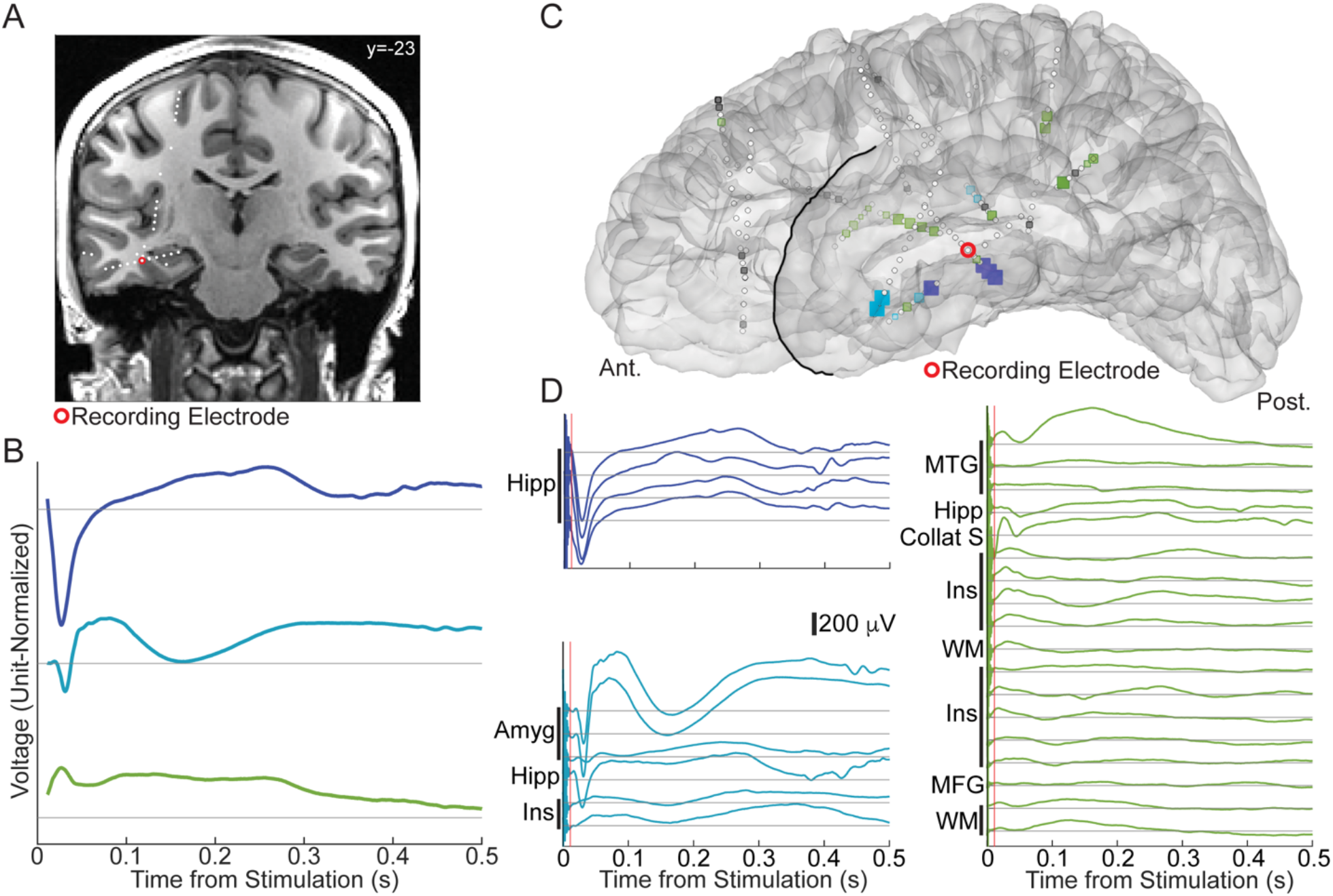
BPCs calculated from CCEP inputs to subject 1, electrode 1 (collateral sulcus). **A**, Coronal slice of subject T1-weighted MRI, depicting location of recording electrode. **B**, Three distinct BPCs were determined using CCEP data from 40 stimulation sites (12 excluded by algorithm), in the 11 to 500 ms interval post-stimulation. **C**, Spatial representation of BPCs by stimulation site (squares) on the brain rendering of subject 1. Colors match BPCs in B; size and color intensity indicate mean SNR across trials at each stimulation site. Stimulation sites are placed at the midpoint of electrode pairs (white circles). Gray indicates stimulation sites discarded by BPC algorithm thresholding. **D**., Mean CCEP from each stimulation site to recording electrode, separated by BPC category and labeled by anatomical location of stimulation site. Amyg = Amygdala, Collat S = Collateral Sulcus, Hipp = Hippocampus, Ins = Insula, MFG = Middle Frontal Gyrus, MTG = Middle Temporal Gyrus, STG = Superior Temporal Gyrus, WM = White Matter.

Between 8 (18%) and 29 (58%) stimulation sites for each recording electrode did not evoke robust responses and did not exceed the threshold set in the algorithm in the calculation of BPC shapes. These stimulation sites were therefore not assigned to any BPCs. Each stimulation site that was assigned to a BPC fit with a degree of robustness quantified by SNR. For example, Figure 2C and 2D demonstrate robust contributions from hippocampal stimulation for the first BPC and from amygdala stimulation for the second BPC.

Similar patterns were observed at other electrodes and subjects. The BPCs tended to be more consistent in shape across recording electrodes within the same subject, though the number of BPCs varied sometimes across recording electrodes within subject due to the absence of a predetermined cluster number.

### Four consensus BPCs across subjects

In individual subjects, the collateral sulcus revealed various contrasting BPC shapes from cortical and subcortical gray stimulation. We tested whether a single set of ‘consensus’ BPCs could map these CCEP shapes across subjects. For this, we first applied singular value decomposition (effectively, principal component analysis without centering) to the set of all 44 BPCs merged across recording electrodes in subjects 1 through 5 (Figure 3A). The first three principal components collectively explained 87% of total variance across BPCs. Next, K-means clustering was performed on the three principal components to identify 4 clusters, their centroids projected back to signal space to yield a set of 4 consensus BPCs. K=4 clusters were chosen based on visual identification of 4 salient clusters of BPCs when projected to the first 3 principal components. However, we note that the dimensionality reduction was unnecessary in this case, as the same 4 clusters were reliably obtained by K-means clustering when any other number of total principal components were kept.

**Figure 3.**
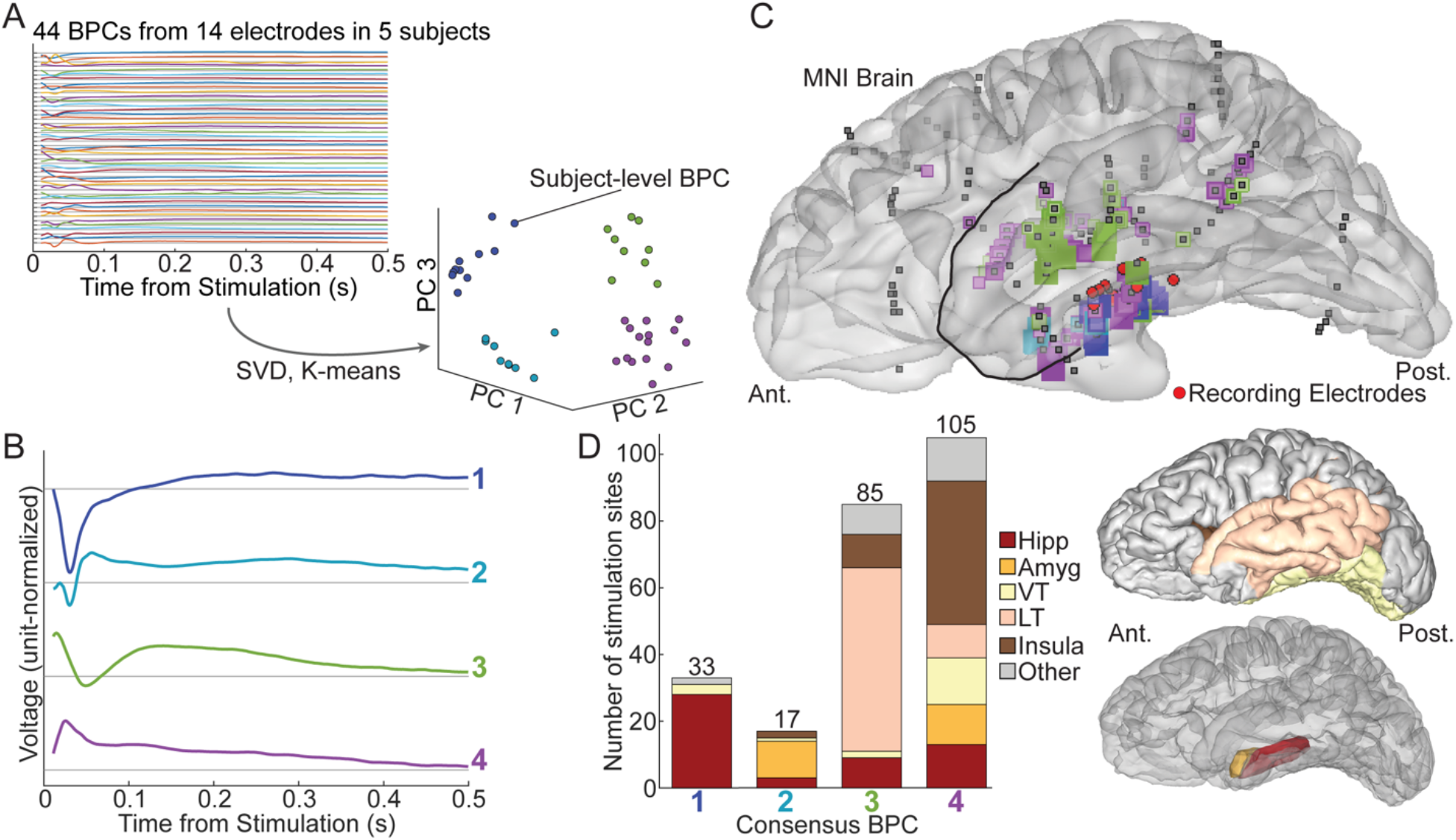
Consensus BPCs and anatomical locations of stimulation sites across 5 subjects. **A**, Consensus BPCs were calculated by applying K-means clustering on all 44 subject-level BPCs from the 14 collateral sulcus electrodes in subjects 1 through 5, after reduction to 3 principal components by singular value decomposition. **B**, Temporal shapes of the 4 consensus BPCs. **C**, Spatial representation of consensus BPCs by stimulation site from subjects 1 through 5, transformed to MNI 152 space, on a standard left hemisphere brain rendering. Stimulation sites with SNR ≥ 1are colored by consensus BPC category, with size and color intensity scaled by subject-level SNR. Gray depicts stimulation sites with SNR < 1 or which were discarded by subject-level BPC algorithm thresholding. **D**, Distribution of stimulation sites across anatomical regions differed significantly between the 4 consensus BPCs (Chi-Square Test of Independence, p = 5.86*10^−43^). Amyg = Amygdala, Hipp = Hippocampus, LT = Lateral Temporal, VT = Ventral Temporal).

Figure 3B shows the consensus BPCs that emerged across the subjects, and their waveform shapes are described in Table 2.

**Table 2.** Origin and shape of collateral sulcus consensus BPCs. Consensus BPCs arose from combinations of electrodes across subjects and demonstrated different waveform patterns. Multiple subject-level BPCs at the same electrode can cluster to the same consensus BPC.

| Consensus BPC ID | Number of subject-level BPCs clustered | Description of consensus BPC |
| --- | --- | --- |
| 1 | 4 from subject 1 (electrodes 1, 2, 3, 4)<br>2 from subject 2 (electrode 1)<br>1 from subject 3 (electrode 1)<br>2 from subject 4 (electrodes 1, 5)<br>2 from subject 5 (electrodes 2, 3) | Single negative peak at 31 ms, followed by a return to a positive steady state over the next 100 ms |
| 2 | 4 from subject 1 (electrodes 1, 2, 3, 4)<br>5 from subject 5 (electrodes 1, 2, 3) | Negative peak at 30 ms, followed by a brief positive peak at 57 ms, and then a wide positive peak centered at 291 ms which then returned gradually to baseline |
| 3 | 1 from subject 2 (electrode 1)<br>1 from subject 3 (electrode 1)<br>7 from subject 4 (electrodes 1, 2, 3, 4, 5) | Positive peak at 15 ms, followed by a minor negative peak at 49 ms, then a wide positive peak at 140 ms which returned gradually to baseline |
| 4 | 4 from subject 1 (electrodes 1, 2, 3, 4)<br>1 from subject 2 (electrode 1)<br>1 from subject 3 (electrode 1)<br>5 from subject 4 (electrodes 1, 2, 3, 4, 5)<br>4 from subject 5 (electrodes 1, 2, 3) | Single positive peak at 26 ms that gradually returned to baseline. |

It is important to note that each consensus BPC was constructed from multiple electrodes across multiple subjects (and thus no single subject drove a particular consensus BPC). In addition, multiple subject-level BPCs with slight differences (e.g., in timing) at one recording electrode could cluster to the same consensus BPC. This allowed for high level consolidation of subject-level BPCs.

### Anatomical distribution of consensus BPCs

Figure 2C demonstrates the anatomical segregation of stimulation sites by BPC cluster in subject 1. For example, stimulation at hippocampal sites often elicited CCEPs assigned to a subject-level BPC exhibiting a single early negative deflection. We tested whether a consistent anatomical pattern would emerge across multiple subjects and electrodes, when grouping stimulation sites by consensus BPC. To answer this question, we tallied the Destrieux atlas labels for all stimulation sites that clustered to each consensus BPC and binned them into 6 major anatomical regions (hippocampus, amygdala, ventral temporal, lateral temporal, insula, and other) (Figure 3D). Stimulation sites that were not assigned to any BPCs (191/585) or were assigned with SNR less than 1 (154/585, lower BPC fit than residual noise) were excluded from this analysis. The anatomical distributions differed significantly across the 4 consensus BPCs (Chi-Square Test of Independence, p = 5.86*10^−43^). As we had observed in single subjects, the majority of stimulation sites that elicited the first consensus BPC were in the hippocampus (28/33, 85%), while the majority of stimulation sites that elicited the second consensus BPC were in the amygdala (11/17, 65%). The third consensus BPC was elicited mostly by stimulation in lateral temporal cortex (55/85, 65%), and the fourth consensus BPC was evoked by stimulation at a broad distribution of sites comprising hippocampus, amygdala, ventral temporal cortex, and insula, but with insula leading in prevalence (43/105, 41%).

To visualize this distribution, electrodes and stimulation sites from all subjects were transformed to MNI 152 coordinates and plotted to a standard left hemisphere MNI 152 pial rendering (Figure 3C) and to axial slices of a standard MNI 152 T1-weighted MRI (Figure 4). On the pial rendering, right hemispheric electrodes were reflected across the midline. It was apparent that although hippocampal stimulation contributed in part to all four consensus BPCs, the finer spatial distribution differed along the longitudinal axis: the second consensus BPC was generally elicited by stimulation more anteriorly in the amygdala/hippocampus, compared to the first consensus BPC. This pattern occurred in both hemispheres and can be seen in greater detail by visualizing stimulation sites on axial and sagittal slices, at the level of the hippocampus, of individual T1 MRI (Figure 7). Cortical and white matter stimulation sites within 6 mm of the pial surface were also rendered onto individual inflated pial renderings (Figure 8).

**Figure 4.**
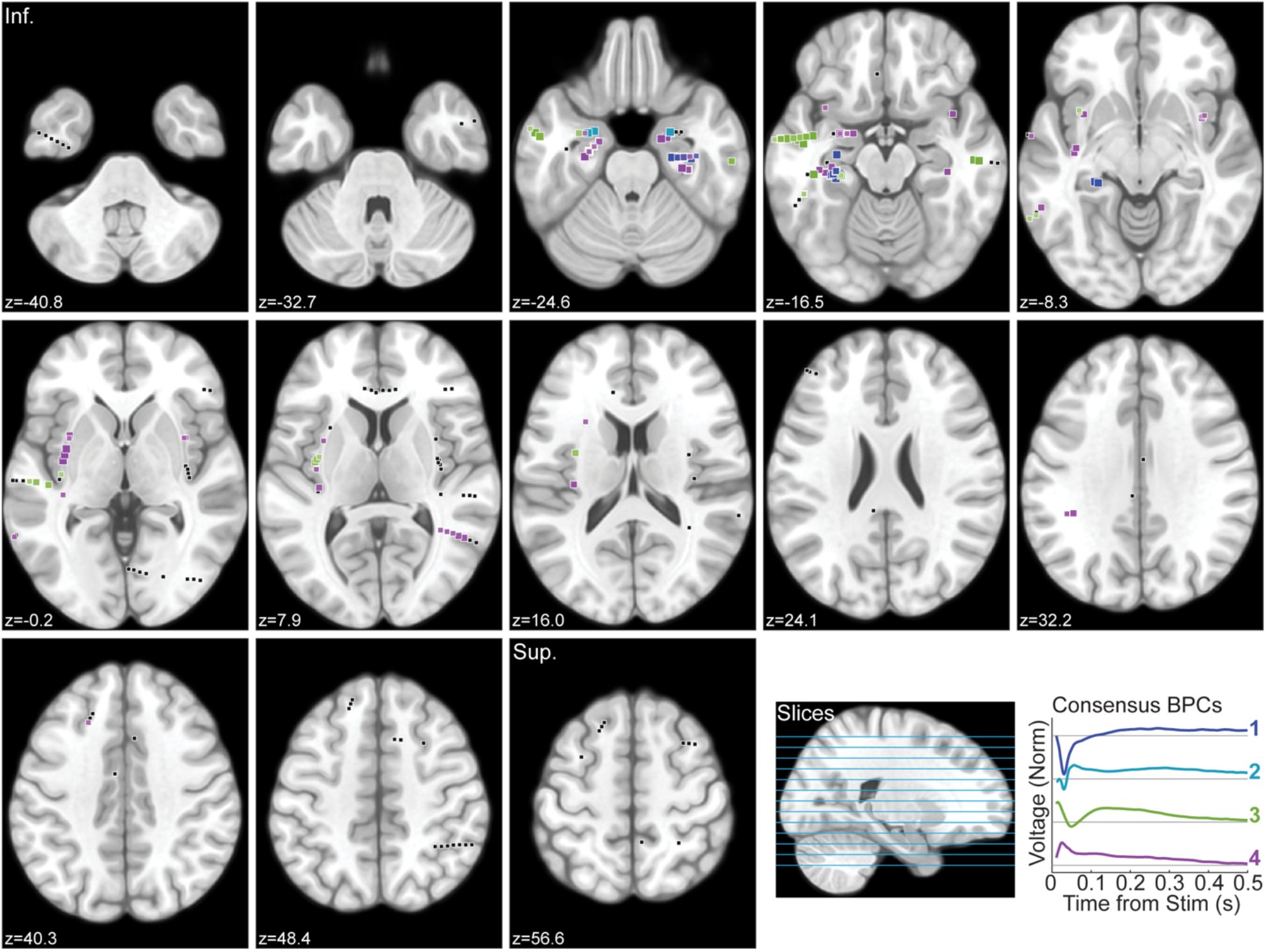
Stimulation sites from all electrodes in subjects 1 through 5 visualized on axial slices of MNI152 T1 MRI. Stimulation sites (squares) were plotted to the nearest slice. Stimulation sites with SNR ≥ 1are colored by consensus BPC category, with size and color intensity scaled by subject-level SNR. Gray depicts stimulation sites with SNR < 1 or which were discarded by subject-level BPC algorithm thresholding. Z-positions of axial slices depicted on sagittal slice in bottom right.

### Spectral and broadband changes

We observed that stimulation of different anatomical clusters evoked differently shaped waveforms in the collateral sulcus. To better understand whether local neuronal populations increase or decrease in activity in response to these various inputs, we calculated time-frequency heatmaps and time-varying broadband estimates (Figure 5). Broadband activity, in particular, has been associated with local neuronal activity (Manning et al., 2009; Miller et al., 2009; Ray and Maunsell, 2011).

**Figure 5.**
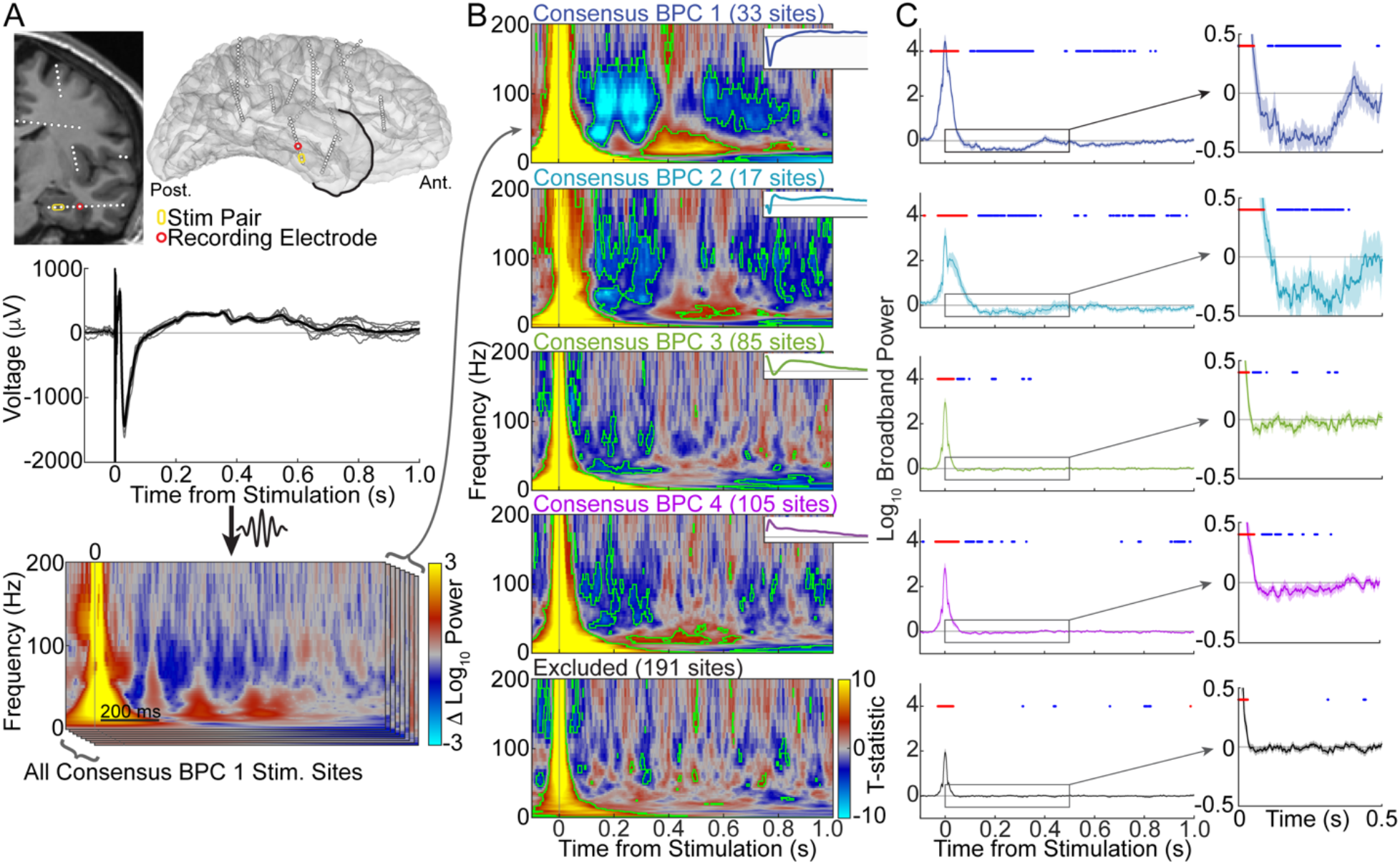
Spectral and broadband power changes of CCEPs across consensus BPC categories. **A**, Example of spectrogram calculation. CCEPs (middle) were measured at subject 2 electrode 1 following stimulation at a hippocampus site assigned to consensus BPC 1 (top). Individual CCEP trials (N=9) in gray and mean across trials in black. An average spectrogram for the stimulation site was calculated by geometrically averaging across all wavelet-transformed trials (bottom: top layer). Average spectrograms were calculated similarly for all other stimulation sites in subjects 1 through 5 assigned to consensus BPC 1 (bottom: stacked layers). **B**, Spectrograms from stimulation sites were combined for each consensus BPC, as well as all sites excluded by the BPC algorithm, by calculating the one-sample t-statistic at each time-frequency bin. Time-frequency bins with significant power change over baseline (corrected for multiple comparisons at FDR < 0.05) are outlined in green. **C**, Time-varying broadband power for the same consensus BPCs in B were calculated by averaging across broadband-filtered CCEPs from stimulation sites. Samples with significant power increase or decrease, relative to baseline, (corrected for multiple comparisons at FDR < 0.05) are marked by red and blue circles, respectively. Inset zooms in on the 0 – 500 ms post-stimulation interval.

We determined the average induced spectrograms for each stimulation site by calculating the geometric mean of spectrograms across trials, after normalizing by baseline. The spectrograms and timevarying broadband estimates from stimulation sites were combined for each consensus BPC, as well as for all stimulation sites algorithmically excluded from BPC assignment (Figures 5B, C). Immediately after stimulation, we observed a rapid broadband increase for all BPCs and excluded stimulation sites up to about 100 ms post-stimulation. Interestingly, the first and second consensus BPCs demonstrated significant decreases in power above 30 Hz, relative to baseline, between 100 and 360 ms post-stimulation (FDR < 0.05, corrected for multiple comparisons). This interval of high frequency power decrease was present to a lesser degree for the third and fourth consensus BPCs, but was not present in excluded sites. In addition, the first and second consensus BPCs induced prolonged intervals of significantly decreased high frequency power from 500 to 1000 ms post-stimulation. Significant increases in power in the 12 to 30 Hz (beta) range were also observed for the first, second, and fourth consensus BPCs between 300 and 650 ms post-stimulation. We note that, despite the large differences in evoked potential responses, relatively similar spectral changes emerged across multiple consensus BPCs.

### Lateral occipito-temporal sulcus inputs

We briefly explored whether similar BPCs would be observed from recording sites in other ventral temporal areas. In our analysis thus far, we have carefully identified electrodes in the collateral sulcus. Similar BPCs in a different VTC region may suggest physiological invariance in the VTC to electrical stimulation inputs. Therefore, we repeated our BPC analysis on inputs to the lateral occipito-temporal sulcus electrodes (see Table 1) in subjects 5 (2 electrodes) and 6 (2 electrodes). Application of the BPC algorithm uncovered sets of 2 to 3 BPCs at each electrode (Figures 6A, C). As with collateral sulcus inputs, we observed consistent BPC shapes and stimulation site clusters between recording electrodes within each of the two subjects.

**Figure 6.**
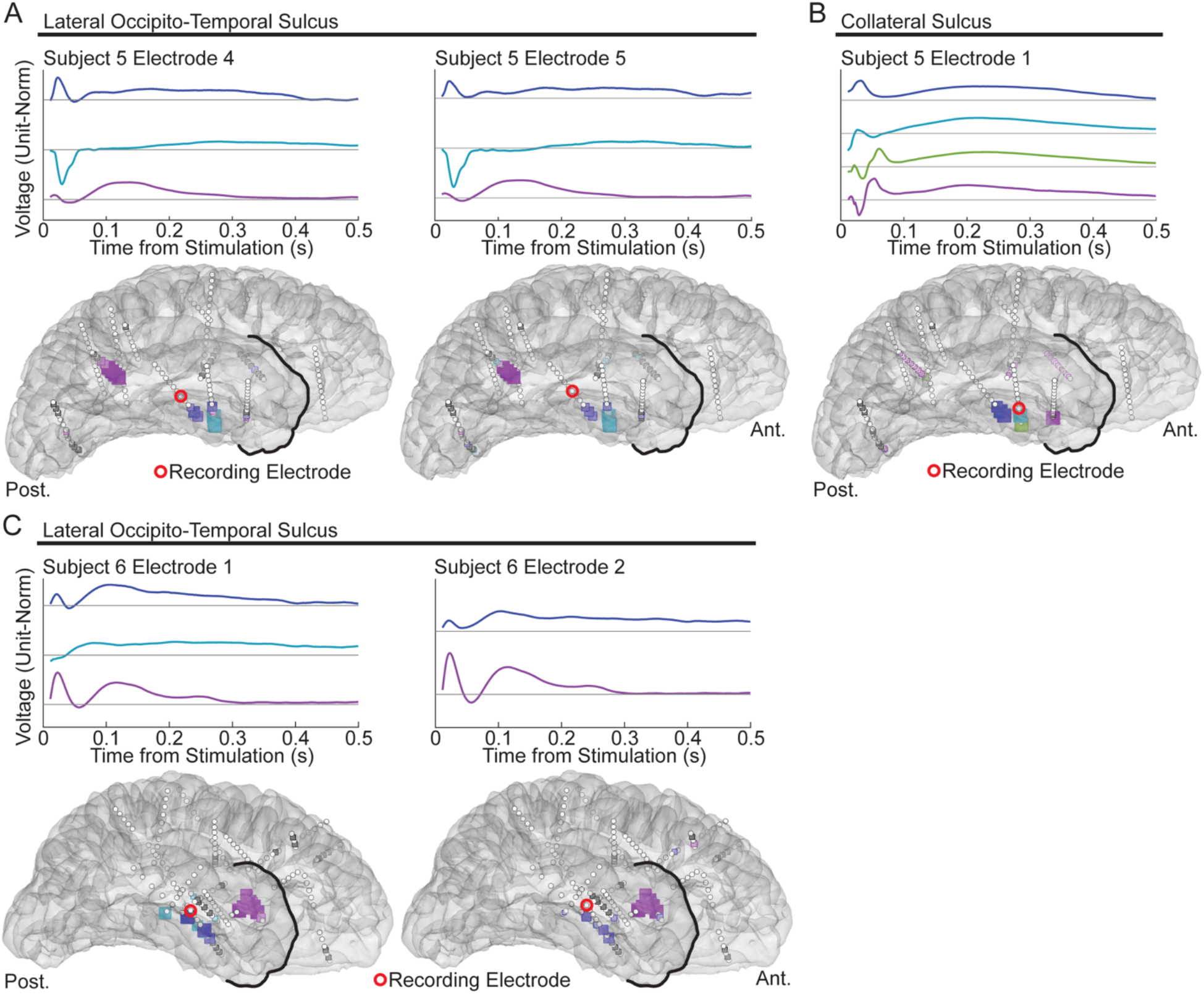
BPCs calculated from CCEP inputs to lateral occipito-temporal sulcus electrodes in subjects 5 and 6. **A**, BPCs calculated from CCEP inputs to subject 5, electrodes 4 and 5 in the lateral occipito-temporal (OT) sulcus. The following are Destrieux atlas labels for stimulation sites that elicited each BPC with SNR > 1. BPC 1 (blue): collateral sulcus and parahippocampal gyrus; BPC 2 (cyan): hippocampus, parahippocampal gyrus, and white matter; BPC 3 (purple): superior temporal sulcus and white matter. **B**, For comparison, BPCs calculated from CCEP inputs to subject 5, electrode 1 in the collateral sulcus. BPC 1 (blue): collateral sulcus and parahippocampal gyrus; BPC 2 (cyan): parahippocampal gyrus; BPC 3 (green): hippocampus; BPC 4 (purple): amygdala. **C**, BPCs calculated from CCEP inputs to subject 6, electrodes 1 and 2 in the lateral occipito-temporal sulcus. BPC 1 (blue): hippocampus; BPC 2 (cyan, electrode 1 only): hippocampus; BPC 3 (purple): insula and white matter. Subject 6 did not have any electrodes in the collateral sulcus.

**Figure 7.**
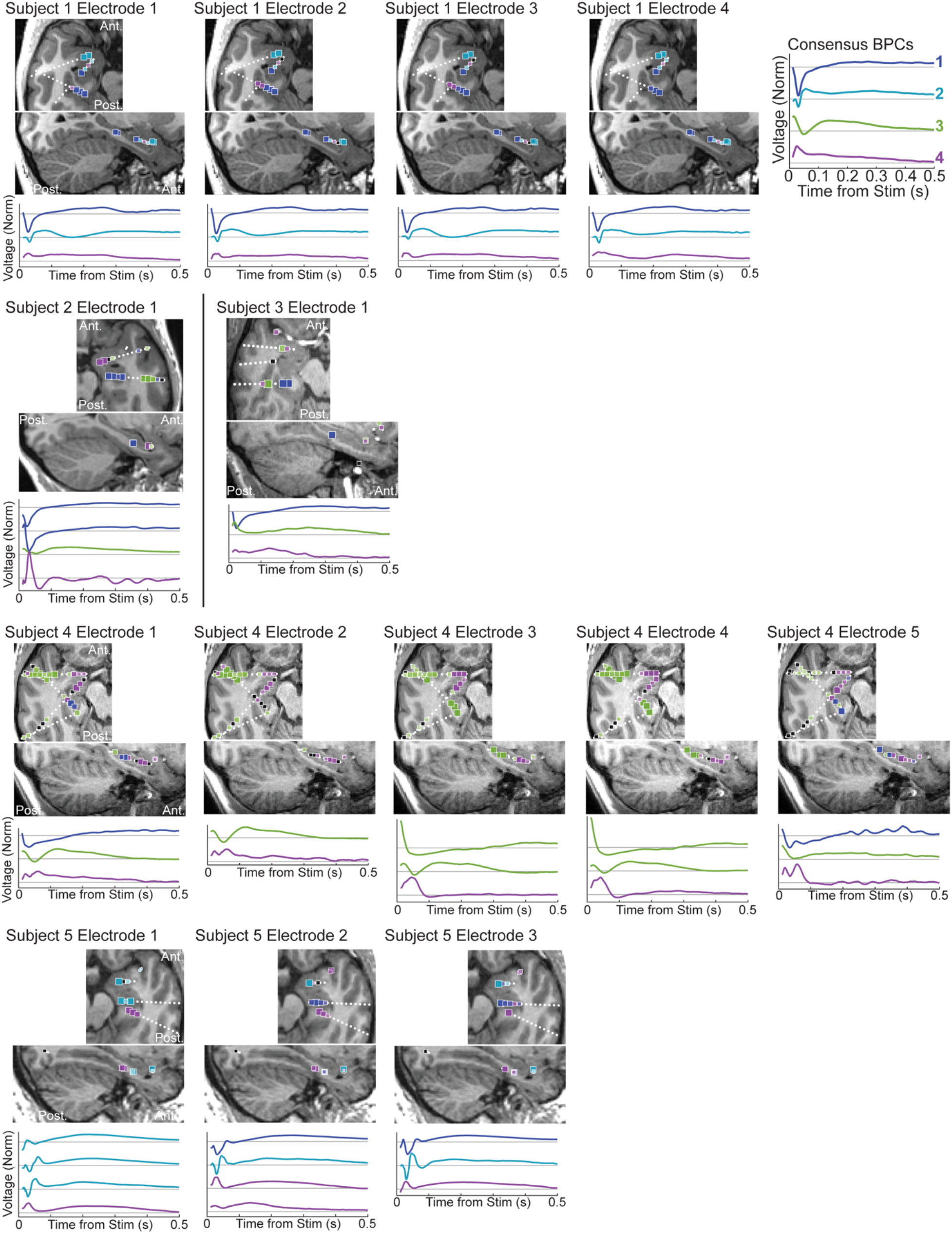
Hippocampal and amygdala stimulation sites. Mid-hippocampal axial and sagittal T1 MRI slices in each subject, with electrodes and stimulation sites within 8 mm distance of each slice visible. Stimulation sites (squares) and subject BPCs were colored according to their assigned consensus BPC for the given recording electrode. Size and color intensity are scaled by subject-level SNR.

**Figure 8.**
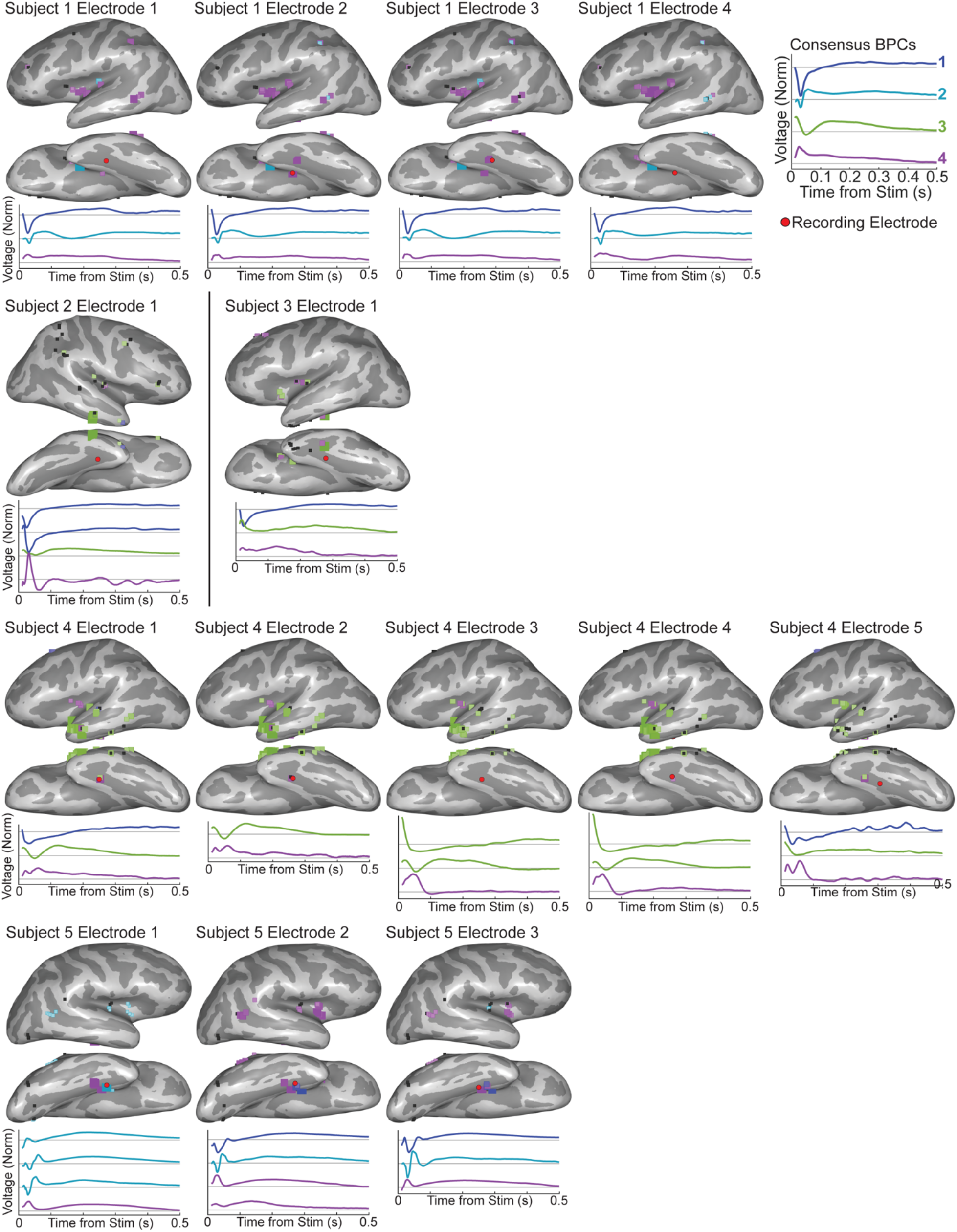
Cortical and white matter stimulation sites within 6 mm of the pial surface in each subjects projected to inflated pial renderings. Stimulation sites (squares) and subject BPCs were colored according to their assigned consensus BPC. Size and color intensity are scaled by subject-level SNR.

Subject 5, who possessed both collateral sulcus electrodes (1, 2, and 3) and lateral occipito-temporal sulcus electrodes (4 and 5), allowed for a direct comparison of input connectivity to the two VTC regions in the same subject. We observed similarities and differences. Stimulation of amygdala evoked a biphasic response in the collateral sulcus (Figure 6B, purple), but no robust response in the lateral occipito-temporal sulcus (Figure 6A). Stimulation of hippocampus also evoked a biphasic response in the collateral sulcus (Figure 6B, green), but a negative-only response in the lateral occipito-temporal sulcus (Figure 6A, cyan). However, stimulation in parahippocampal gyrus produced similar positive deflections in both VTC regions (Figures 6A, B, blue). These differences underscore the heterogeneity of connectivity to different regions of the VTC.

## DISCUSSION

In this study, we characterized single pulse electrical stimulation-driven inputs to the collateral sulcus and the lateral occipito-temporal sulcus within the VTC. We discovered that CCEP shapes varied substantially by stimulation site, and that those shapes broadly segregated by the anatomical regions stimulated. The unique CCEP input shapes were summarized at each recording electrode by sets of 2 to 4 waveforms (BPCs), and again across electrodes and subjects by a single set of 4 consensus BPC waveforms. Stimulation produced a period of high frequency power decrease between 100 and 360 ms post-stimulation regardless of BPC category, though the effect was strongest with stimulation sites that evoked the first and second consensus BPCs, from predominantly hippocampal and amygdala stimulation.

Across subjects, stimulation in the hippocampus tended to elicit a prominent early negative deflection, while stimulation in lateral temporal cortex sites and insula elicited an early positive deflection. The primary advantage in keeping the recording site fixed is that differences in CCEP shape reflect different physiologic connectivity to the same population of measured neurons. Early studies using the current source density method in cat visual cortex have demonstrated that local field potential (LFP) polarity recorded from microelectrodes largely depends on synaptic configuration in local neuronal populations (Eccles, 1951; Mitzdorf and Singer, 1978). For example, pronounced negative deflections in surface LFP often reflect dipolar current sinks produced by synchronized excitatory postsynaptic potentials (EPSPs) at apical dendrites of superficial pyramidal cells (Mitzdorf, 1985). In contrast, positive deflections in surface LFP may reflect superficial inhibitory postsynaptic potentials (IPSPs) or EPSPs at deeper layers.

The information contained in BPC waveforms may thus help to reveal the type of pathway that connects the stimulated and measured populations. For instance, lateral temporal cortex inputs to the collateral sulcus contained a long positive deflection. Lateral temporal cortex inputs to the fusiform face area, adjacent to the collateral sulcus, have been investigated with fMRI and iEEG. These studies have shown that the superior temporal sulcus is involved in processing dynamic facial features such as gaze (Gobbini and Haxby, 2007; Pitcher and Ungerleider, 2021; Babo-Rebelo et al., 2022), despite an absence of direct connections to the fusiform face area (Ethofer et al., 2011; Gschwind et al., 2012; Pyles et al., 2013). It has been proposed therefore, that the ventral visual stream may be indirectly connected to the lateral temporal cortex through the parietal cortex or through the inferior temporal cortex via short range U-fibers (Babo-Rebelo et al., 2022). These models of indirect connectivity are consistent with the long time course of the lateral temporal input to the collateral sulcus, uncovered by our BPC analysis.

Different waveforms may also provide estimations on how likely the post-synaptic neuronal population is to fire from afferent stimulation, as they may contain information pertaining to the coherence of afferent EPSPs, as well as physical distance between synapse and soma (Mitzdorf, 1985). CCEP waveforms may also comprise action potentials and neuronal spikes elicited by electrical stimulation in both orthodromic and antidromic directions (Fuortes et al., 1957; Phillips, 1959). Finally, large deflections in CCEPs might reflect physiologic low-frequency rhythms that have been phase reset and synchronized by stimulation (Nakae et al., 2018). Further analysis of temporal dynamics using BPCs may help to elucidate the extent of contribution from these various factors.

The results presented here indicate that CCEP responses may be more complex than previously assumed. Many ECoG-CCEP studies have focused on the earliest negative potential in the initial 50 ms, termed the N1, which is often followed by a later negative deflection termed the N2 (Matsumoto et al., 2004, 2007; Keller et al., 2014; Araki et al., 2015; Krieg, 2017; van Blooijs et al., 2018; Kundu et al., 2020; Silverstein et al., 2020). The BPCs we derived from sEEG recordings contained reliable waveforms that differed from this pattern. The absence of this N1/N2 pattern is likely attributable, in part, to electrical field differences from measurements at various cortical depths using sEEG electrodes, compared to measurements at the cortical surface with ECoG. Our sEEG electrodes were positioned closer to the gray-white matter boundary than the pial surface. At this depth, the N1/N2 peaks would reverse: that is, signals measured deeper to a synaptic potential generator would result in inverted polarity from the surface potential (Mitzdorf and Singer, 1978). Indeed, the two positive peaks in the third consensus BPC (Figure 3B), measured at deeper cortical layers (e.g., see location of electrode in Figure 2A) might represent inverted N1 and N2 peaks if recorded at the cortical surface instead.

Our data showed differences in BPC waveforms recorded in the collateral sulcus compared to the lateral occipito-temporal sulcus, the two regions being separated by the mid-fusiform sulcus (MFS). The VTC has been shown to be sharply delineated by the MFS into medial and lateral components with differing functional and anatomical features (Grill-Spector and Weiner, 2014). Functionally, cortex medial to the MFS is preferentially activated by visual stimuli that are peripheral, inanimate, large, and which contain places; while cortex lateral to the MFS is preferentially activated by stimuli that are foveal, animate, small, and which contain faces (Nasr et al., 2011; Konkle and Oliva, 2012; Weiner et al., 2014). Structurally, the MFS marks a medial-lateral separation in cytoarchitectonics and white matter connectivity (Saygin et al., 2012; Weiner et al., 2014; Gomez et al., 2015). The place-selective region in the collateral sulcus further differs in white matter connectivity to face-, body-, and word-selective regions lateral to the MFS, in terms of both fascicle and endpoint connectivity profiles (Kubota et al., 2022). Such anatomical boundaries are therefore important to consider when comparing CCEP waveforms across different recording sites.

CCEPs permit directed measurements of connectivity between stimulated and recording sites and can be regarded as a form of effective connectivity, comparable to other measures of brain connectivity. A recent study found that connectivity networks based on CCEP N1 amplitude demonstrate significant similarity to connectivity networks constructed by voltage correlation, coherence, and diffusion tensor imaging (DTI) (Crocker et al., 2021). Specifically, the authors noted that at local distances to the stimulation site, CCEP connectivity most resembled (functional) correlation connectivity, and at longer distances, CCEP connectivity most resembled (structural) DTI connectivity.

Spectral analysis of inputs to the collateral sulcus consistently revealed significantly decreased high frequency broadband power, above 30 Hz and between 100 ms and 360 ms post-stimulation. Broadband power has been correlated with underlying neuronal (input) spiking activity (Manning et al., 2009; Miller et al., 2009; Ray and Maunsell, 2011). As such, our results suggest inhibited neuronal activity following single pulse electrical stimulation of connected sites. Previous studies have similarly found that single pulse or low frequency electrical stimulation in gray matter tended to produce long lasting neuronal inhibition (Alarcón et al., 2012; Westin et al., 2018; Mohan et al., 2020). Plausible physiologic explanations for this include hyperpolarization mediated by GABAergic interneurons or slow after-hyperpolarization (Toprani and Durand, 2013), and in fact, inhibition in monkey visual cortex induced by electrical stimulation in lateral geniculate nucleus could be disrupted with GABA antagonists (Logothetis et al., 2010). Recurrent feedback inhibition involving such GABAergic interneurons is key to cortical circuits as a means of gain control following strong stimulation (Markram et al., 2004; Douglas and Martin, 2007; Ozeki et al., 2009).

The recordings in our analysis were conducted in patients affected by intractable epilepsy, with large variations in age, sex, and epilepsy etiology. Electrodes were placed in regions indicated by the clinical team, which constrained electrode selection in the VTC across subjects. This may have contributed to between-subject variability in the BPC shapes calculated. In addition, both the collateral sulcus and lateral occipitotemporal sulcus span several centimeters longitudinally, so there may have been different types of inputs to recording electrodes placed in different parts of the sulci (Weiner et al., 2018; Kubota et al., 2022). This variability was mitigated by the fact that we did not include electrodes close to the anterior or posterior collateral transverse sulci, as they were labelled separately by the Freesurfer segmentation. Disparity in electrode placement also meant that CCEPs may have been affected to varying degrees by volume conduction when stimulating in nearby areas (Prime et al., 2020). Finally, more detailed anatomical segmentations may further delineate differences in BPC categorization. The waveform differences along the longitudinal axis of the hippocampus may have arisen from different hippocampal subfields (Strange et al., 2014). Despite these potential sources of variability, consensus BPC waveforms correlated relatively well to broad anatomical regions across multiple recording electrodes and subjects.

In conclusion, we found that single pulse electrical stimulation inputs to two anatomical areas in the VTC could be described by characteristic time-varying waveforms. Some of these shapes were distinct from the archetypal “N1” and “N2” peaks. The shapes segregated by stimulation location in cortical and limbic areas with robustness across subjects and may contain valuable information for mapping synaptic physiology at the recording site. High frequency power decreases post-stimulation spanned across stimulation categories and may be characteristic of stimulation-induced inhibition of neuronal activity.

## Acknowledgements

We are grateful for the participation of the patients in this study, and for the assistance of Cindy Nelson, Karla Crockett, and other staff at Saint Mary Hospital, Mayo Clinic, Rochester, MN. Research reported in this publication was supported by the National Institute of Mental Health of the National Institutes of Health under Award Number R01MH122258. The content is solely the responsibility of the authors and does not represent the official views of the National Institutes of Health.

## REFERENCES

Alarcón G, Martinez J, Kerai SV, Lacruz ME, Quiroga RQ, Selway RP, Richardson MP, García Seoane JJ, Valentín A (2012) In vivo neuronal firing patterns during human epileptiform discharges replicated by electrical stimulation. Clinical Neurophysiology 123:1736–1744.

Araki K, Terada K, Usui K, Usui N, Araki Y, Baba K, Matsuda K, Tottori T, Inoue Y (2015) Bidirectional neural connectivity between basal temporal and posterior language areas in humans. Clinical Neurophysiology 126:682–688.

Babo-Rebelo M, Puce A, Bullock D, Hugueville L, Pestilli F, Adam C, Lehongre K, Lambrecq V, Dinkelacker V, George N (2022) Visual Information Routes in the Posterior Dorsal and Ventral Face Network Studied with Intracranial Neurophysiology and White Matter Tract Endpoints. Cerebral Cortex 32:342–366.

Benjamini Y, Yekutieli D (2001) The Control of the False Discovery Rate in Multiple Testing under Dependency. The Annals of Statistics 29:1165–1188.

Bressler SL (1995) Large-scale cortical networks and cognition. Brain Research Reviews 20:288–304.

Crocker B, Ostrowski L, Williams ZM, Dougherty DD, Eskandar EN, Widge AS, Chu CJ, Cash SS, Paulk AC (2021) Local and distant responses to single pulse electrical stimulation reflect different forms of connectivity. NeuroImage 237:118094.

Dale AM, Fischl B, Sereno MI (1999) Cortical surface-based analysis: I. Segmentation and surface reconstruction. Neuroimage 9:179–194.

Destrieux C, Fischl B, Dale A, Halgren E (2010) Automatic parcellation of human cortical gyri and sulci using standard anatomical nomenclature. NeuroImage 53:1–15.

Douglas RJ, Martin KAC (2007) Mapping the Matrix: The Ways of Neocortex. Neuron 56:226–238.

Eccles JC (1951) Interpretation of action potentials evoked in the cerebral cortex. Electroencephalography and Clinical Neurophysiology 3:449–464.

Edelman GM, Mountcastle VB (1978) The mindful brain: Cortical organization and the group-selective theory of higher brain function. The mindful brain: Cortical organization and the group-selective theory of higher brain function: 100–100.

Epstein R, Kanwisher N (1998) A cortical representation of the local visual environment. Nature 392:598–601.

Ethofer T, Gschwind M, Vuilleumier P (2011) Processing social aspects of human gaze: a combined fMRI-DTI study. Neuroimage 55:411–419.

Felleman DJ, Van Essen DC (1991) Distributed Hierarchical Processing in the Primate Cerebral Cortex. Cerebral Cortex 1:1–47.

Friston KJ (1994) Functional and effective connectivity in neuroimaging: A synthesis. Human Brain Mapping 2:56–78.

Fuortes MGF, Frank K, Becker MC (1957) Steps in the production of motoneuron spikes. The Journal of general physiology 40:735–752.

Gobbini MI, Haxby JV (2007) Neural systems for recognition of familiar faces. Neuropsychologia 45:32–41.

Goldman-Rakic PS (1988) Topography of cognition: Parallel distributed networks in primate association cortex. Annual Review of Neuroscience 11:137–156.

Gomez J, Pestilli F, Witthoft N, Golarai G, Liberman A, Poltoratski S, Yoon J, Grill-Spector K (2015) Functionally Defined White Matter Reveals Segregated Pathways in Human Ventral Temporal Cortex Associated with Category-Specific Processing. Neuron 85:216–227.

Grill-Spector K, Weiner KS (2014) The functional architecture of the ventral temporal cortex and its role in categorization. Nat Rev Neurosci 15:536–548.

Gschwind M, Pourtois G, Schwartz S, Van De Ville D, Vuilleumier P (2012) White-matter connectivity between face-responsive regions in the human brain. Cereb Cortex 22:1564–1576.

Hermes D, Miller KJ, Noordmans HJ, Vansteensel MJ, Ramsey NF (2010) Automated electrocorticographic electrode localization on individually rendered brain surfaces. Journal of neuroscience methods 185:293–298.

Holdgraf C et al. (2019) iEEG-BIDS, extending the Brain Imaging Data Structure specification to human intracranial electrophysiology. Sci Data 6:102.

Huang H, Valencia GO, Hermes D, Miller KJ (2021) A canonical visualization tool for SEEG electrodes. In: 2021 43rd Annual International Conference of the IEEE Engineering in Medicine Biology Society (EMBC), pp 6175–6178.

Kanwisher N, McDermott J, Chun MM (1997) The Fusiform Face Area: A Module in Human Extrastriate Cortex Specialized for Face Perception. J Neurosci 17:4302.

Kay KN, Yeatman JD (2017) Bottom-up and top-down computations in word- and face-selective cortex. eLife 6:e22341.

Keller CJ, Honey CJ, Mégevand P, Entz L, Ulbert I, Mehta AD (2014) Mapping human brain networks with cortico-cortical evoked potentials. Philosophical Transactions of the Royal Society B: Biological Sciences 369:20130528.

Konkle T, Oliva A (2012) A real-world size organization of object responses in occipitotemporal cortex. Neuron 74:1114–1124.

Krieg J (2017) Discrimination of a medial functional module within the temporal lobe using an effective connectivity model: A CCEP study. :13.

Kubota E, Grotheer M, Finzi D, Natu VS, Gomez J, Grill-Spector K (2022) White matter connections of high-level visual areas predict cytoarchitecture better than category-selectivity in childhood, but not adulthood. Cerebral Cortex:bhac221.

Kundu B, Davis TS, Philip B, Smith EH, Arain A, Peters A, Newman B, Butson CR, Rolston JD (2020) A systematic exploration of parameters affecting evoked intracranial potentials in patients with epilepsy. Brain Stimulation 13:1232–1244.

Lacruz ME, García Seoane JJ, Valentin A, Selway R, Alarcón G (2007) Frontal and temporal functional connections of the living human brain. European Journal of Neuroscience 26:1357–1370.

Logothetis NK, Augath M, Murayama Y, Rauch A, Sultan F, Goense J, Oeltermann A, Merkle H (2010) The effects of electrical microstimulation on cortical signal propagation. Nat Neurosci 13:1283–1291.

Manning JR, Jacobs J, Fried I, Kahana MJ (2009) Broadband Shifts in Local Field Potential Power Spectra Are Correlated with Single-Neuron Spiking in Humans. J Neurosci 29:13613–13620.

Markov NT, Vezoli J, Chameau P, Falchier A, Quilodran R, Huissoud C, Lamy C, Misery P, Giroud P, Ullman S, Barone P, Dehay C, Knoblauch K, Kennedy H (2014) Anatomy of hierarchy: Feedforward and feedback pathways in macaque visual cortex. Journal of Comparative Neurology 522:225–259.

Markram H, Toledo-Rodriguez M, Wang Y, Gupta A, Silberberg G, Wu C (2004) Interneurons of the neocortical inhibitory system. Nat Rev Neurosci 5:793–807.

Matsumoto R, Nair DR, LaPresto E, Bingaman W, Shibasaki H, Lüders HO (2007) Functional connectivity in human cortical motor system: a cortico-cortical evoked potential study. Brain 130:181–197.

Matsumoto R, Nair DR, LaPresto E, Najm I, Bingaman W, Shibasaki H, Lüders HO (2004) Functional connectivity in the human language system: a cortico-cortical evoked potential study. Brain 127:2316–2330.

McManus JNJ, Li W, Gilbert CD (2011) Adaptive shape processing in primary visual cortex. PNAS 108:9739–9746.

Mesulam M-M (1990) Large-scale neurocognitive networks and distributed processing for attention, language, and memory. Annals of Neurology 28:597–613.

Mewett DT, Nazeran H, Reynolds KJ (2001) Removing power line noise from recorded EMG. In: 2001 Conference Proceedings of the 23rd Annual International Conference of the IEEE Engineering in Medicine and Biology Society, pp 2190–2193 vol.3.

Miller KJ, Müller K-R, Hermes D (2021) Basis profile curve identification to understand electrical stimulation effects in human brain networks van Vugt MK, ed. PLoS Comput Biol 17:e1008710.

Miller KJ, Sorensen LB, Ojemann JG, den Nijs M (2009) Power-Law Scaling in the Brain Surface Electric Potential. PLOS Computational Biology 5:e1000609.

Mitzdorf U (1985) Current source-density method and application in cat cerebral cortex: investigation of evoked potentials and EEG phenomena. Physiological Reviews 65:37–100.

Mitzdorf U, Singer W (1978) Prominent excitatory pathways in the cat visual cortex (A 17 and A 18): A current source density analysis of electrically evoked potentials. Exp Brain Res 33:371–394.

Mohan UR, Watrous AJ, Miller JF, Lega BC, Sperling MR, Worrell GA, Gross RE, Zaghloul KA, Jobst BC, Davis KA, Sheth SA, Stein JM, Das SR, Gorniak R, Wanda PA, Rizzuto DS, Kahana MJ, Jacobs J (2020) The effects of direct brain stimulation in humans depend on frequency, amplitude, and white-matter proximity. Brain Stimulation 13:1183–1195.

Moran J, Desimone R (1985) Selective attention gates visual processing in the extrastriate cortex. Science 229:782–784.

Motter BC (1993) Focal attention produces spatially selective processing in visual cortical areas V1, V2, and V4 in the presence of competing stimuli. J Neurophysiol 70:909–919.

Nakae T, Matsumoto R, Togo M, Takeyama H, Kobayashi K, Shimotake A, Matsuhashi M, Yamao Y, Kikuchi T, Yoshida K, Kunieda T, Ikeda A, Miyamoto S (2018) S128. Oscillatory responses evoked by single-pulse electrical stimulation in human cerebral cortex –A Cortico-Cortical Evoked Potential (CCEP) study. Clinical Neurophysiology 129:e189–e190.

Nasr S, Liu N, Devaney KJ, Yue X, Rajimehr R, Ungerleider LG, Tootell RB (2011) Scene-selective cortical regions in human and nonhuman primates. Journal of Neuroscience 31:13771–13785.

Ozeki H, Finn IM, Schaffer ES, Miller KD, Ferster D (2009) Inhibitory stabilization of the cortical network underlies visual surround suppression. Neuron 62:578–592.

Penny WD, Friston KJ, Ashburner JT, Kiebel SJ, Nichols TE (2011) Statistical parametric mapping: the analysis of functional brain images. Elsevier.

Phillips CG (1959) Actions of antidromic pyramidal volleys on single Betz cells in the cat. Quarterly Journal of Experimental Physiology and Cognate Medical Sciences: Translation and Integration 44:1–25.

Pitcher D, Ungerleider LG (2021) Evidence for a Third Visual Pathway Specialized for Social Perception. Trends Cogn Sci 25:100–110.

Prime D, Woolfe M, O’Keefe S, Rowlands D, Dionisio S (2020) Quantifying volume conducted potential using stimulation artefact in cortico-cortical evoked potentials. Journal of Neuroscience Methods 337:108639.

Pyles JA, Verstynen TD, Schneider W, Tarr MJ (2013) Explicating the Face Perception Network with White Matter Connectivity. PLOS ONE 8:e61611.

Ray S, Maunsell JHR (2011) Different Origins of Gamma Rhythm and High-Gamma Activity in Macaque Visual Cortex. PLOS Biology 9:e1000610.

Saygin ZM, Osher DE, Koldewyn K, Reynolds G, Gabrieli JD, Saxe RR (2012) Anatomical connectivity patterns predict face selectivity in the fusiform gyrus. Nature neuroscience 15:321–327.

Schlack A, Albright TD (2007) Remembering visual motion: neural correlates of associative plasticity and motion recall in cortical area MT. Neuron 53:881–890.

Silverstein BH, Asano E, Sugiura A, Sonoda M, Lee M-H, Jeong J-W (2020) Dynamic tractography: Integrating cortico-cortical evoked potentials and diffusion imaging. NeuroImage 215:116763.

Sporns O, Chialvo DR, Kaiser M, Hilgetag CC (2004) Organization, development and function of complex brain networks. Trends in Cognitive Sciences 8:418–425.

Strange BA, Witter MP, Lein ES, Moser EI (2014) Functional organization of the hippocampal longitudinal axis. Nat Rev Neurosci 15:655–669.

Toprani S, Durand DM (2013) Long-lasting hyperpolarization underlies seizure reduction by low frequency deep brain electrical stimulation. The Journal of physiology 591:5765–5790.

van Blooijs D, Leijten FSS, van Rijen PC, Meijer HGE, Huiskamp GJM (2018) Evoked directional network characteristics of epileptogenic tissue derived from single pulse electrical stimulation. Human Brain Mapping 39:4611–4622.

Weiner KS, Barnett MA, Witthoft N, Golarai G, Stigliani A, Kay KN, Gomez J, Natu VS, Amunts K, Zilles K, Grill-Spector K (2018) Defining the most probable location of the parahippocampal place area using cortex-based alignment and cross-validation. NeuroImage 170:373–384.

Weiner KS, Golarai G, Caspers J, Chuapoco MR, Mohlberg H, Zilles K, Amunts K, Grill-Spector K (2014) The mid-fusiform sulcus: A landmark identifying both cytoarchitectonic and functional divisions of human ventral temporal cortex. NeuroImage 84:453–465.

Westin K, Lundstrom B, Cooray G (2018) F05. Neurophysiological effects of chronic subthreshold cortical stimulation for focal epilepsy: Spike and spontaneous ECoG activity. Clinical Neurophysiology 129:e67–e68.

